# Tissue nanotransfection-mediated induction of neurogenic programs promotes myoprotective responses in denervated skeletal muscle

**DOI:** 10.64898/2026.07.10.737742

**Authors:** Ana I. Salazar-Puerta, Sara Kheirkhah, Jordan T. Moore, Carlos A. Vasquez-Martinez, Carolina Velasquez-Quintero, Hallie Harris, Manami Fukuda, Megumi E. Fukuda, Jonathan P. Stranan, Fangli Zhao, Kavya Dathathreya, Jared Albert, Prameela Bobbili, Charles D. Wendt, Jonathan Winograd, Ian L. Valerio, Candice Askwith, Amy M. Moore, W. David Arnold, Daniel Gallego Perez

**Affiliations:** Department of Biomedical Engineering, The Ohio State University, Columbus, OH, USA; Department of Plastic and Reconstructive Surgery, The Ohio State University, Columbus, OH, USA; Department of Neuroscience, The Ohio State University, Columbus, OH, USA; Department of Neurology, The Ohio State University Wexner Medical Center, Columbus, OH, USA; Plastic and Reconstructive Surgery, Massachusetts General Hospital, Boston, MA, USA; Physical Medicine and Rehabilitation, University of Missouri, Columbia, MO, USA; Department of Surgery, The Ohio State University, Columbus, OH, USA

**Keywords:** Peripheral nerve injury, muscle protection, cell reprogramming, neuroengineering

## Abstract

Peripheral nerve injuries often result in prolonged skeletal muscle denervation, leading to progressive atrophy, fibrosis, neuromuscular instability, and loss of regenerative capacity before axons can reinnervate distal targets. Here, we developed a non-viral strategy using tissue nanotransfection (TNT) to deliver the neurogenic transcription factor cocktail *Ascl1*, *Brn2*, and *Myt1l* (*ABM*) directly to denervated skeletal muscle. *In vitro*, *ABM*-transfected myoblasts sustained expression of the reprogramming factors, acquired neuron-like morphologies, upregulated neuronal markers including Tuj1, Map2, and Syp, and exhibited electrophysiological properties consistent with membrane excitability. RNA sequencing confirmed broad activation of neurogenic transcriptional programs, with enrichment of pathways associated with neuronal fate commitment, neuron differentiation, axon guidance, synaptogenesis, and developmental signaling. In a mouse model of sciatic nerve transection, TNT enabled localized *ABM* expression in denervated gastrocnemius muscle. *ABM*-TNT treatment accelerated resolution of denervation-associated fibrillation potentials and showed trends toward improved twitch and tetanic torque, compound muscle action potential amplitudes, and muscle mass preservation. Transcriptomic profiling of treated muscles 5 weeks after injury revealed distinct gene expression programs enriched for muscle regeneration, neuromuscular organization, trophic support, extracellular matrix remodeling, angiogenesis, myogenesis, and metabolic adaptation. Network analyses further identified activation of neurogenic regulators, neurotrophic signaling, and vascular-support pathways. These findings establish TNT-mediated *ABM* delivery as a non-viral platform for inducing neurogenic and myoprotective programs in denervated muscle, suggesting a potential strategy to preserve muscle viability during the prolonged interval required for peripheral nerve regeneration.

## INTRODUCTION

Peripheral nerve injuries, often resulting from traumatic disruption of peripheral nerves in both civilian and military settings, frequently lead to substantial sensorimotor impairment, long-term disability, neuropathic pain, and incomplete functional recovery[1]. Following severe injury or transection, the distal nerve segment undergoes Wallerian degeneration, motor endplates lose neural input, and the denervated muscle rapidly progresses toward eventual irreversible atrophy and fibrosis[2]. Since axonal regeneration occurs at only 1–2 mm/day, months may pass before regenerating axons reach their distal targets and initiate reinnervation. During this prolonged interval, denervated skeletal muscle undergo a rapid and progressive degenerative process driven by the loss of electrophysiological and biochemical communication between the nerve and muscle[3–5]. With prolonged denervation, muscle fibers are continuously lost through apoptosis, satellite cell responsiveness declines, and the vascular network degenerates, collectively reducing the muscle’s capacity for functional recovery even once axonal regrowth eventually reaches the affected tissue[6].

Human studies indicate that denervated muscle and motor endplates retain the capacity for successful reinnervation for approximately 18–24 months, and beyond this period, functional recovery drops significantly. Fibrosis and scar tissue begin forming around 12 months and become irreversible, and by two years, extreme muscle atrophy and structural degeneration preclude restoration of function[7–9]. Under normal physiological conditions, Schwann cells play a central role in supporting axonal regeneration; however, this supportive phenotype is both temporally limited and spatially constrained to regions near the injury site. Furthermore, for injuries involving gaps larger than 5–50 mm, graft-based repair strategies continue to fall short of providing complete functional recovery[10]. These limitations indicate that promoting axonal regeneration alone is insufficient and that preserving a muscle environment that remains viable and receptive to reinnervation is critical for improving long-term functional outcomes[11].

Previous studies investigated the ability of stem cell-derived neurons to establish synaptic connections with muscle[12, 13]. However, these approaches require extensive *ex vivo* manipulation, including the generation and differentiation of neural progenitors prior to transplantation, and are associated with an inherent risk of tumorigenicity. Pluripotent stem cell-derived and neural progenitor cell-based grafts may retain undifferentiated or highly plastic cells that have been shown to form teratomas or malignant tumors in preclinical models, representing a major safety barrier to clinical implementation[14, 15]. To overcome these limitations, direct transdifferentiation strategies were developed to bypass induced pluripotent stem cell (iPSC) intermediates and generate functional neurons directly from adult somatic cells[16, 17]. This approach offers a faster and potentially safer alternative to stem cell-based therapies. Nevertheless, most current direct-conversion protocols rely heavily on viral vectors, which present significant barriers to clinical translation due to safety concerns, including immunogenicity, insertional mutagenesis, and limit capacity for repeat dosing[18]. These limitations underscore the need for clinically translatable delivery platforms capable of inducing neuronal reprogramming without the safety concerns associated with viral vectors.

Our group[19–21] and others[22–24] have previously demonstrated that the transcription factors (TFs) *Ascl1, Brn2, and Myt1l*, referred as *ABM*, can directly convert somatic cells, including fibroblasts, glial cells, hepatocytes, into neurons[25, 26]. Successful direct neuronal reprogramming requires TFs capable of activating neuronal gene networks while suppressing the donor-cell identity. The *ABM* cocktail fulfills these requirements through complementary mechanisms: *Ascl1* acts as a pioneer factor that opens closed chromatin and initiates neuronal gene activation[17], *Brn2* reinforces neuronal identity by suppression non-neuronal programs[27], and *Myt1l* promotes neuronal maturation and synaptic function[28].

To enable the delivery of these TFs into solid tissues, our group developed Tissue Nano-Transfection (TNT), a non-viral, nanochannel-based platform that uses brief electrical pulses to drive genetic cargo into solid tissues *in situ*, using the host as a bioreactor for cellular reprogramming[19, 29, 30]. We hypothesized that TNT-mediated delivery of the ABM cocktail to skeletal muscle would induce neurogenic programs in resident muscle cells and promote the emergence of neuron-like phenotypes capable of supporting muscle integrity during denervation. Skeletal muscle represents an attractive target for *in situ* neuronal induction because it contains resident progenitor populations, including myoblasts that retain cellular plasticity and localize to regions where motor endplates degenerate following nerve injury[31]. Moreover, the muscle microenvironment may support the induction and persistence of neurogenic programs, providing a potential strategy to preserve tissue viability when axonal regeneration alone is insufficient to restore function.[32–34].

In this study, we investigated whether TNT-mediated delivery of *ABM* could induce neurogenic reprogramming responses that support muscle preservation during prolonged denervation. We first characterize *ABM*-driven cellular and transcriptional changes *in vitro* over 7-21 days and performed RNA sequencing (RNA-seq) to evaluate activation of neurogenic gene programs in myoblasts. We then applied TNT-mediated *ABM* delivery to denervated muscle *in vivo* and assessed functional, histological, and transcriptional outcomes 5 weeks after injury. Transcriptomic analysis revealed sustained activation of pathways associated with muscle preservation, structural remodeling, and cellular metabolism, together with engagement of neurogenic and vasculogenic signaling networks. These molecular changes were accompanied by electrophysiological evidence of improved neuromuscular stability, suggesting that TNT-mediated *ABM* delivery promotes a tissue state conducive to preserving muscle integrity during denervation. Collectively, these findings support the potential of non-viral reprogramming-based strategies to maintain muscle viability and function during the prolonged interval required for reinnervation.

## RESULTS

### Myoblasts support prolonged ABM expression and activation of neurogenic programs

Previous studies from our group and others, have demonstrated efficient *ABM*-mediated conversion of fibroblasts into induced neurons[19, 20, 22, 35]. However, because skeletal muscle is the target tissue for TNT-mediated delivery *in vivo*, it is important to determine whether muscle-derived cells exhibit a similar response to neurogenic transcription factors. Notably, skeletal muscle myoblasts possess chromatin features that are permissive to *Ascl1*-mediated neuronal induction[28], suggesting that they may be particularly amenable to activation of neurogenic gene programs. Therefore, we evaluated the response of murine myoblasts to *ABM* delivery *in vitro*. Given the relevance of myoblasts to the denervated muscle environment, we next evaluated whether *ABM* could induce activation of neurogenic programs in murine myoblasts. To this end, C2C12 cells, an immortalized mouse myoblast cell line derived from the thigh muscle of C3H mice, were electrotransfected with expression plasmids encoding for *ABM*, and sham-transfected cells (vector only) were included as controls **(Figure 1A)**. Gene expression analysis confirmed robust upregulation of *Ascl1, Brn2, Myt1l* transcripts in *ABM*-treated myoblasts relative to sham 24 hours post-transfection **(Figure 1B)**. To verify co-expression of the TF cocktail, each plasmid was engineered to express a distinct fluorescent reporter: *Ascl1-*green fluorescent protein (GFP), *Brn2*-red fluorescent protein (RFP), and *Myt1l*-cyan fluorescent protein (CFP). Cells positive for all three fluorophores demonstrated simultaneous expression of the complete *ABM* cocktail 72 hours post-transfection. As expected, sham-transfected cells expressed only the GFP reporter present in the backbone plasmid **(Figure 1C)**. Finally, expression of the *ABM* factors was detected in myoblast cultures 7 days post-transfection, followed by a progressive decline at 14 and 21 days, while remaining elevated relative to sham-transfected controls **(Figure 1D)**.

**Figure 1.**
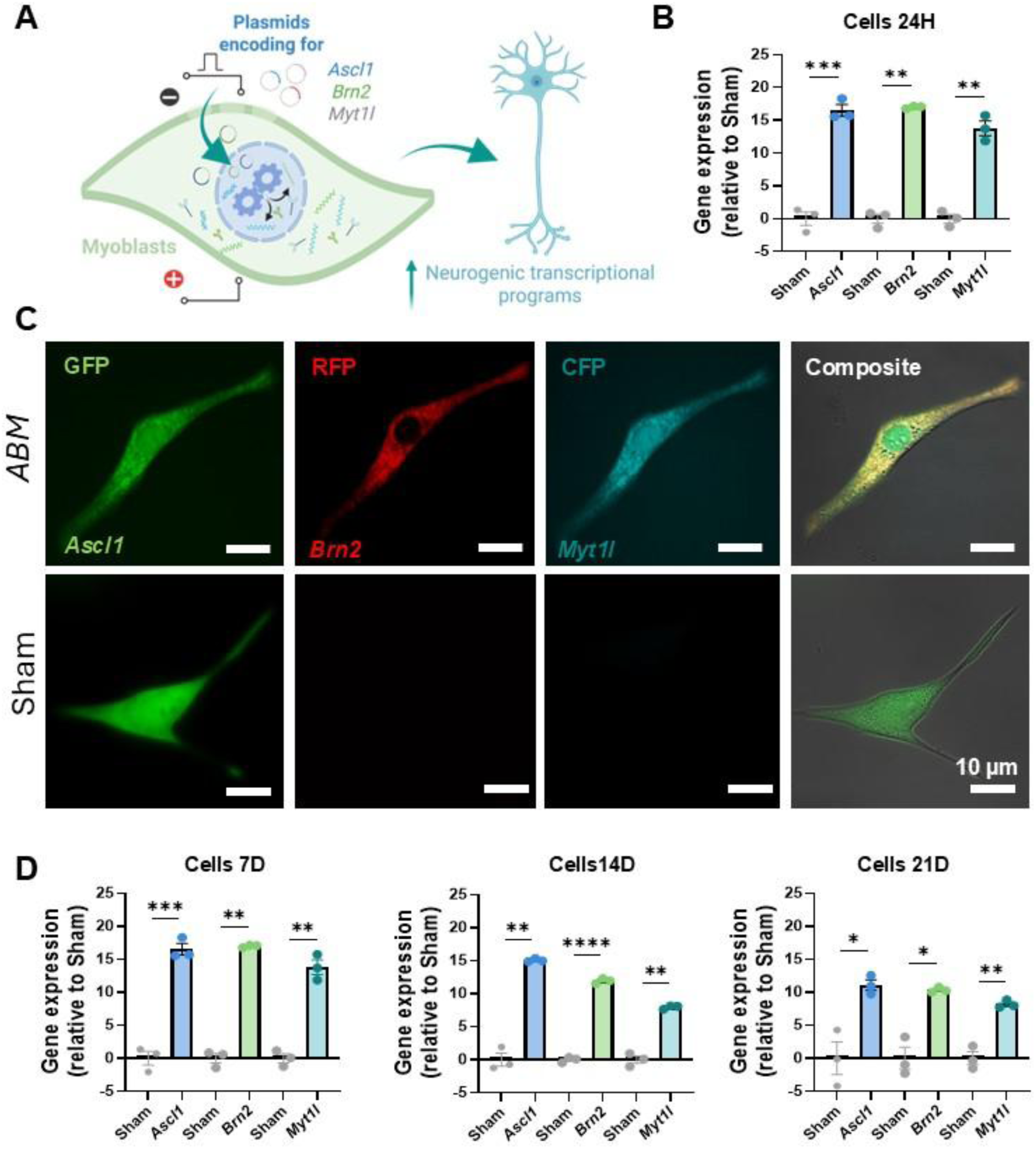
Electrotransfection mediates sustained expression of ABM neurogenic transcription factors in myoblasts. **(A)** Schematic illustrating delivery of the neurogenic transcription factors *Ascl1*, *Brn2*, and *Myt1l* (*ABM*) to myoblasts via electrotransfection, resulting in activation of neurogenic transcriptional programs and acquisition of neuronal-like features. **(B)** qRT-PCR analysis demonstrating robust expression of *Ascl1*, *Brn2*, and *Myt1l* in the *ABM*-transfected myoblasts compared to sham controls 24 hours post-transfection. **(C)** Representative fluorescence images showing co-expressing Ascl1-GFP, Brn2-RFP, and Myt1l-CFP 72 hours after *ABM* transfection compared to sham-transfected myoblasts expressing only the GFP reporter encoded by the control backbone plasmid. **(D)** Longitudinal gene expression analysis of *Ascl1*, *Brn2*, and *Myt1l* using qRT-PCR at 7, 14, and 21 days post-transfection relative to sham-treated cells. (n=3) All error bars are shown as SEM. *p-value<0.05, One-way ANOVA. Created in BioRender. Salazar, A. (2026).

To determine whether *ABM* delivery induced neurogenic responses over time, we assessed neuronal marker expression, morphological changes, and electrophysiological activity. We quantified the expression of neuronal markers including Tubulin Beta 3 Class III (*Tuj1*), microtubule associated protein 2 (*Map2*), and Synaptophysin (*Syp*). Because cultures were not sorted to enrich for fully reprogrammed cells, digital PCR (dPCR) was used to enable sensitive and absolute quantification of transcript abundance across the entire cell population. dPCR analysis revealed a marked increase in *Tuj1* transcript copy number in *ABM*-treated cells compared to sham controls as early as 7 days post-transfection **(Figure 2A)**. By day 14, *ABM-*treated myoblasts exhibited significantly higher expression of *Tuj1*, *Map2*, and *Syp* relative to sham controls **(Figure 2B)**. Finally, *Map2* and *Syp* expression remained detectable in the *ABM* group compared to sham-transfected cells after 21 days post-transfection, indicating sustained activation of neurogenic gene programs following *ABM* delivery **(Figure 2C)**. Consistent with these molecular changes, *ABM*-treated myoblasts underwent morphological remodeling. As early as 7 days post-transfection, cells exhibited bipolar and multipolar morphologies with elongated neurite-like projections, features that were absent in sham-transfected cultures **(Figure 2D)**. To assess whether these cells acquired electrophysiological properties associated with excitable cells, patch-clamp recordings were performed at day 21 on *ABM*-treated myoblasts. Depolarizing voltage steps evoked transient inward currents consistent with activation of voltage-gated sodium (Na+) channels, followed by sustained outward currents consistent with voltage-gated potassium (K+) currents **(Figure 2E)**. In addition, current injection elicited single action potentials in cells exhibiting bipolar morphology, indicating the acquisition of membrane excitability **(Figure 2F)**. Together, these findings demonstrate that myoblasts support prolonged *ABM* expression and undergo molecular, morphological, and electrophysiological changes consistent with partial acquisition of neuronal programs.

**Figure 2.**
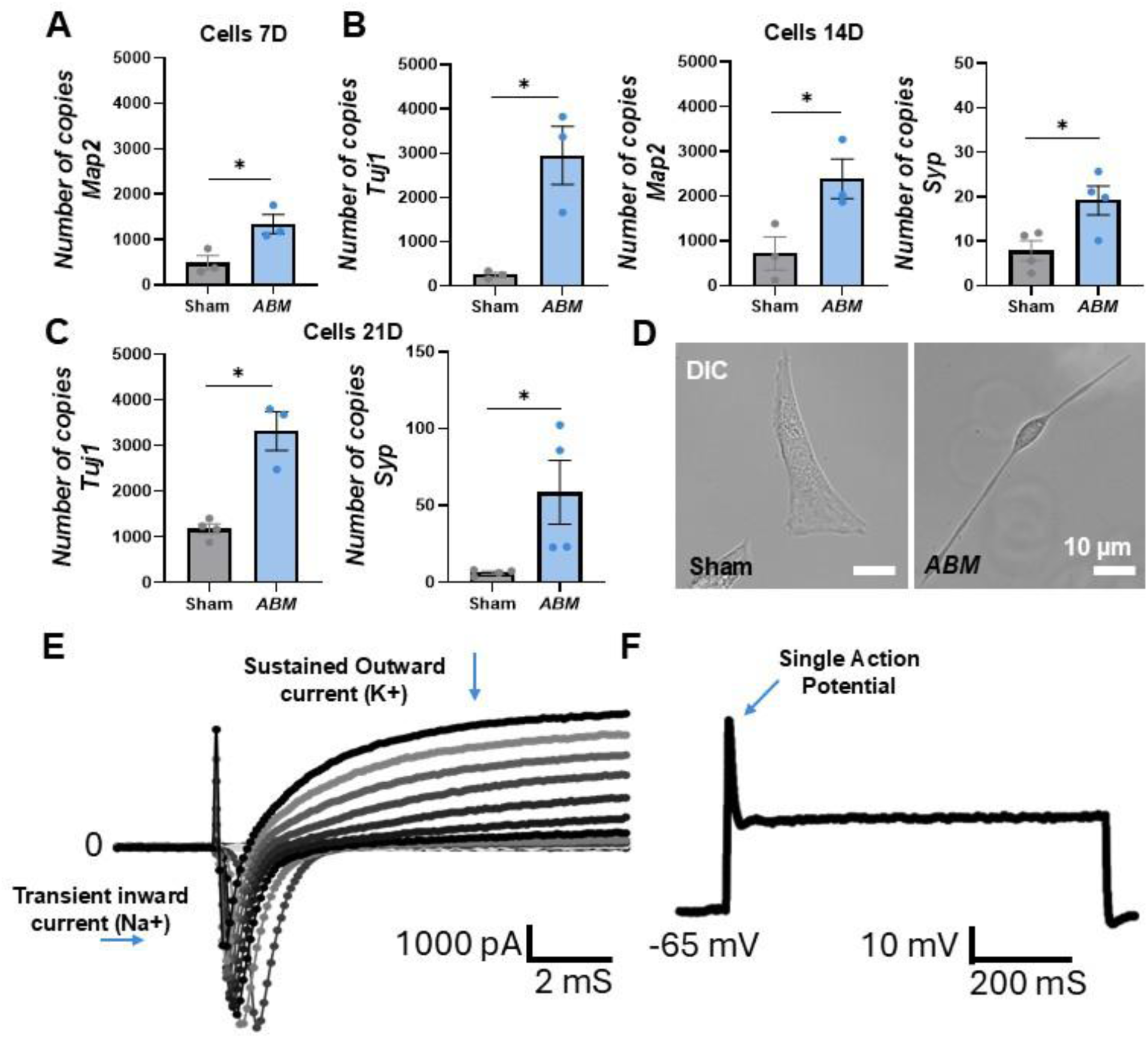
*ABM* delivery induces neurogenic molecular, morphological, and electrophysiological changes in myoblasts. dPCR analysis revealed increased absolute copy numbers of neuronal markers, including **(A)** *Tuj1* at 7 days post-transfection, **(B)** *Tuj1*, *Map2*, and *Syp* at 14 days post-transfection, and **(C)** *Map2* and *Syp* at 21 days post-transfection compared to sham-transfected cells. **(D)** Representative images demonstrating neural-like morphological changes, including elongated neurite-like projections and bipolar cell morphologies in *ABM*-transfected cells relative to sham at 7 days post-transfection. **(E)** Representative whole-cell voltage-clamp recording from *ABM*-treated bipolar myoblast. Depolarizing voltage steps elicited a rapidly activating transient inward current followed by sustained outward currents, consistent with activation of voltage-gated sodium and potassium channels. **(F)** Representative recording demonstrating an evoked action potential in an *ABM*-treated bipolar myoblast in response to a 300pA current injection. Recordings are representative of at least three cells. (n=3-4) All error bars are shown as SEM. *p-value<0.05, t-test.

### ABM induces neurogenic transcriptional programs in myoblasts

To comprehensively characterize the molecular changes induced by *ABM* delivery, whole-transcriptome RNA sequencing (RNA-seq) was performed on myoblast cultures collected 14 days after transfection. Sham-transfected (vector-only) and non-transfected cultures were included as controls **(Figure S1A)**. Differentially expressed gene (DEG) analysis identified 1,590 significantly upregulated and 581 downregulated genes in the *ABM* group relative to sham-transfected controls **(Figure 3A)**. Comparison of *ABM*-treated cells with non-transfected controls similarly revealed substantial transcriptional remodeling with 410 upregulated and 595 downregulated genes (**Figure S1B**). Gene Ontology (GO) analysis of DEGs demonstrated significant enrichment of biological processes (BP) associated with neuronal lineage acquisition, including neuron fate commitment, positive regulation of neuron differentiation, and neuron projection guidance in *ABM*-treated myoblasts relative to sham controls **(Figure 3B)**. Consistent with these findings, DEG clustering revealed enrichment of neurogenic gene signatures in the *ABM* group relative to sham-transfected **(Figure 3C and Figure S1C)** and non-transfected controls **(Figure S1E)**, further supporting a distinct neurogenic transcriptional state. Notably, the *ABM* group showed increased expression of *Ascl1, Pou3f2 (Brn2), and Myt1l* compared to sham, reflecting successful and prolonged expression of the reprogramming transcription factors by day 14 and activation of downstream neuronal programs. In addition, genes associated with neuronal differentiation and maturation were upregulated in the *ABM* group, including *Ndrg4, Necab1, Tubb3, Gprin3, Dlg2,* and *Ctxn3.* These genes are consistent with neuronal differentiation[36], and with the acquisition of neuronal structural, functional, and signaling properties indicative of neuronal identity[37, 38] and maturation[39, 40] **(Figure 3C-D)**. Correspondingly, heatmap analyses demonstrated clear segregation of *ABM*-treated from both sham-transfected **(Figure S1D)** and non-transfected controls **(Figure S1F)**. To gain insight into the biological pathways underlying these changes, gene set enrichment analysis (GSEA) was performed. Compared with sham-transfected controls, *ABM*-treated cultures exhibited significant enrichment of Hedgehog signaling, Notch signaling, and epithelial-to-mesenchymal transition (EMT) pathways. These pathways have established roles in neurogenesis[41], and in cellular plasticity and remodeling processes that facilitate lineage transitions[42, 43].

**Figure 3.**
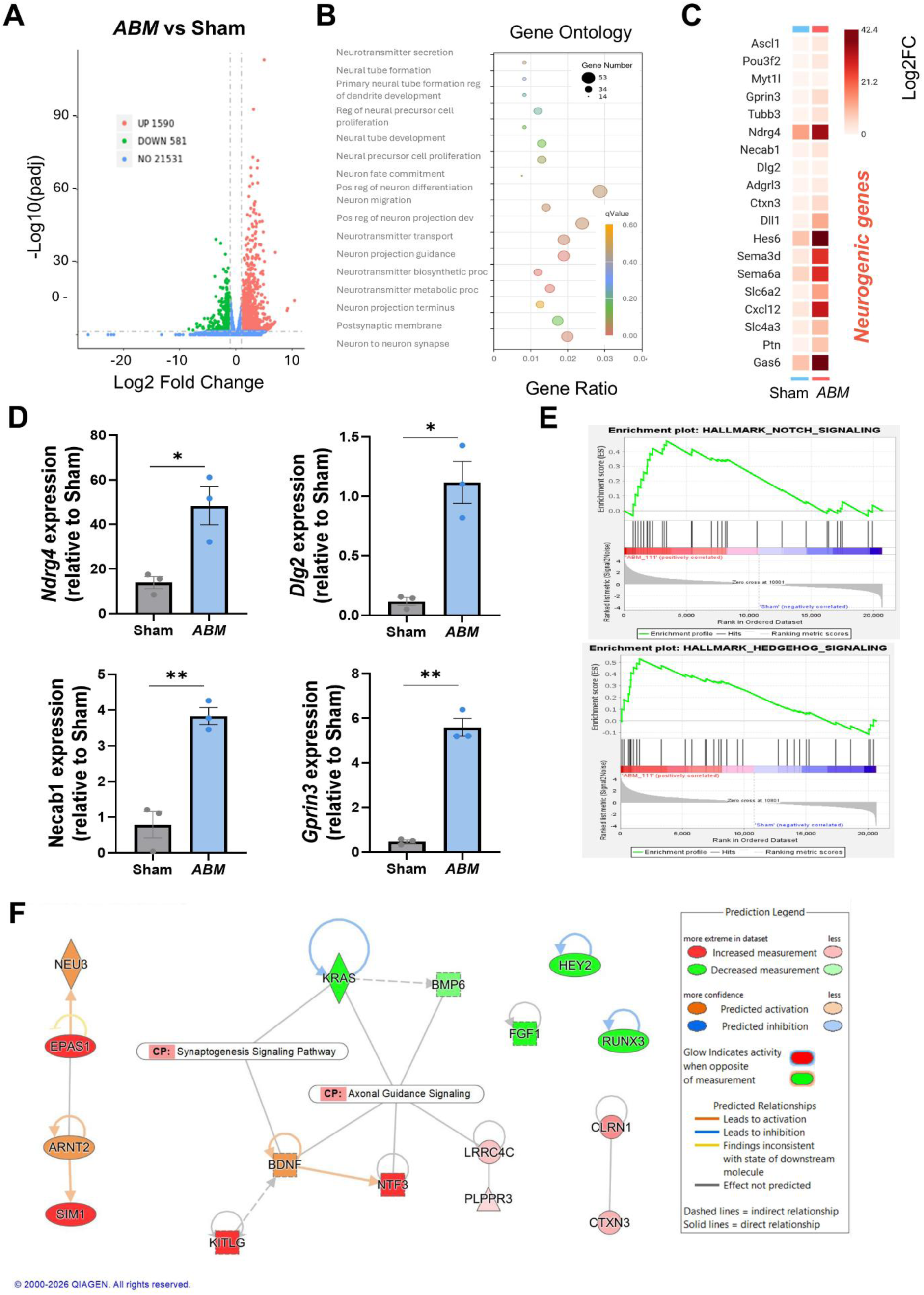
*ABM* drives neurogenic transcriptional programs in myoblasts. RNA-seq analysis was performed on myoblast cultures 14 days after electrotransfection with plasmids encoding *ABM*, compared to sham-transfected cells. **(A)** Volcano plots depicting differentially expressed genes (DEGs) in *ABM*-treated cells compared to sham. Upregulated genes are shown in red, downregulated genes in green, and non-significant genes in blue. **(B)** Gene Ontology (GO) analysis demonstrating enrichment of biological processes related to neuron fate commitment, positive regulation of neuron differentiation, neuron projection guidance in *ABM* group compared to sham-transfected cells. **(C-D)** DEG clustering heatmaps and bar plots showing the enrichment of neurogenic gene signatures associated with neuronal differentiation and maturation were upregulated in the *ABM* group relative to sham **(E)** Gene set enrichment analysis (GSEA) demonstrating enrichment of developmental signaling pathways, including Notch and Hedgehog signaling, on *ABM-*treated cells. **(F)** Ingenuity pathway analysis (IPA) network illustrating activation of neurogenic signaling nodes and concurrent predicted inhibition of muscle-associated pathways in *ABM*-treated myoblasts. IPA network legend shown on the right side. Genes with Log2FC ≥∣1∣ and adjusted p-value ≤ 0.05 were considered significantly differentially expressed.

To further examine pathway-level changes, Ingenuity Pathway Analysis (IPA) was performed. Network analysis revealed enrichment of transcriptional programs associated with neuronal development and maturation in *ABM*-treated cells. Notably, the transcription factors SIM1 and ARNT2, both implicated in neuronal fate specification and maturation[44], emerged as central nodes within the predicted regulatory network. IPA additionally identified activation of components of the synaptogenesis signaling pathway, including BDNF and KITLG, which are associated with neuronal survival, neurite extension, and synaptic development[45]. In contrast, BMP6, a regulator associated with progenitor maintenance and mesenchymal or muscle-related programs[46], was predicted to be inhibited, suggesting suppression of non-neuronal lineage programs. Consistent with this shift, genes involved in axonal and synaptic development, including NTF3 and LRRC4C, converged within the axonal guidance signaling pathway, reinforcing the progression toward a neuronal phenotype induced by *ABM*[47, 48] **(Figure 3F)**. Similarly, when comparing *ABM*-treated cultures to non-transfected cells, NTF3 seems to be upregulated in this pathway, supporting neuronal survival, differentiation, and maintenance[49]. Additional activation of neuronal regulatory modules included TARDBP and growth factors such as PTN and FGF21, which have been implicated in neuronal survival, migration and neurite outgrowth[50, 51], further supporting engagement of neuronal identity programs. Conversely, MRTFA, a transcriptional coactivator involved in skeletal muscle development[52], was predicted to be inhibited, suggesting attenuation of the myogenic program **(Figure S1G)**. Collectively, these results indicate that the *ABM* delivery induces extensive transcriptional remodeling in myoblasts characterized by activation of neurogenic gene networks, enrichment of neuronal differentiation and maturation pathways, and suppression of myogenic regulatory programs. These data support the acquisition of a neurogenic transcriptional state and neuronal-like molecular features following *ABM* treatment.

### Tissue Nano-Transfection-mediated delivery of ABM induces myoprotective responses in denervated muscle

TNT is an emerging non-viral delivery platform that enables highly efficient, localized, and rapid (millisecond-scale) transfer of molecular cargo into solid tissues[29, 30, 53–56]. TNT facilitates the delivery of charged molecular cargo, such as plasmid DNA, through silicon microneedles integrated with nanochannels. Upon application of a focused electric field, transient membrane permeabilization and electrophoretic transport allow cargo to enter target cells. The structure and integrity of the TNT device were confirmed by scanning electron microscopy (SEM) **(Figure 4A)**.

**Figure 4.**
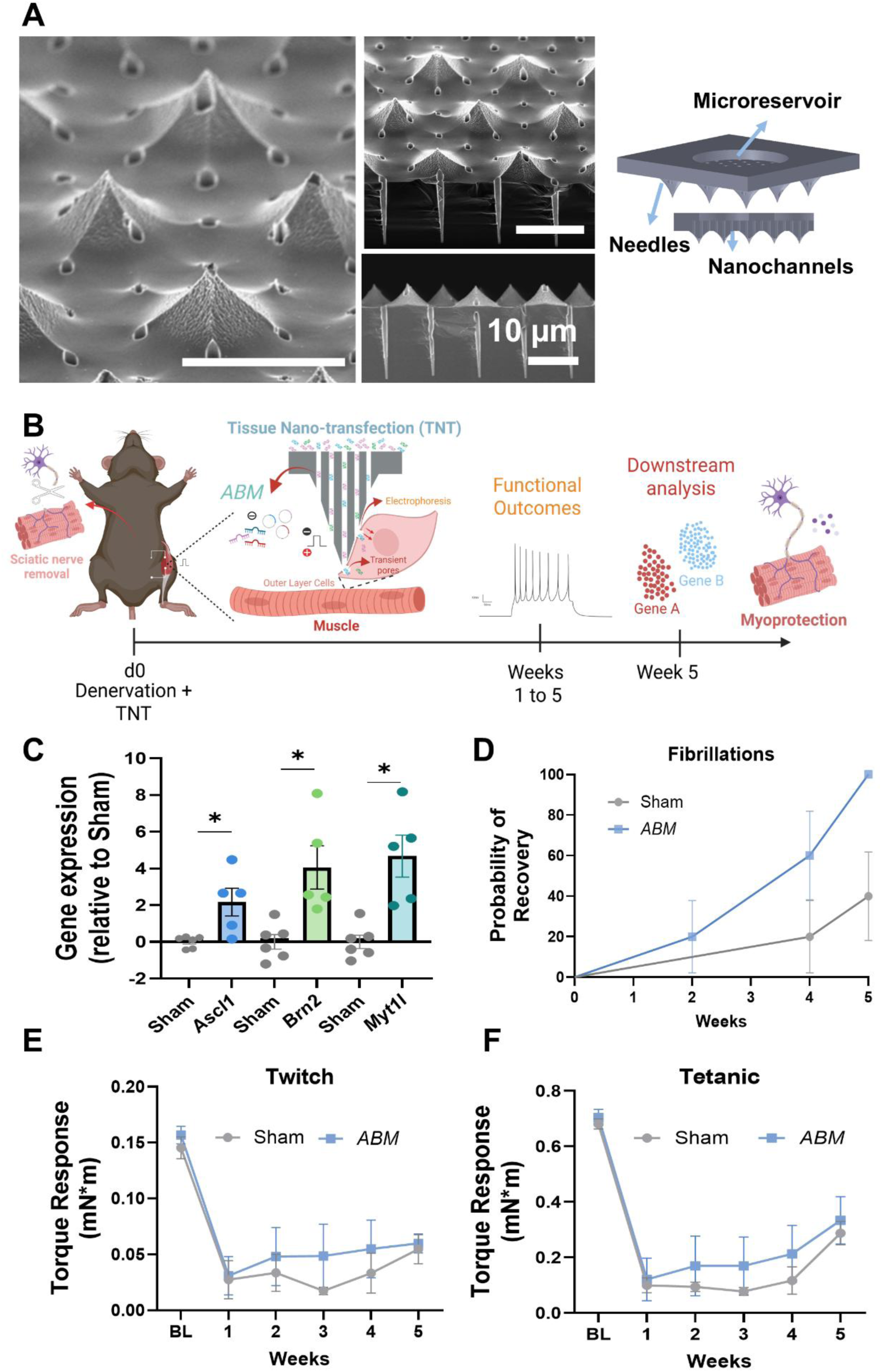
TNT-mediated delivery of *ABM* promotes myoprotective responses in denervated skeletal muscle: **(A)** Schematic of the tissue nano-transfection (TNT) device illustrating the microreservoir, microneedles, and nanochannel architecture. Representative scanning electron microscopy (SEM) images show top and cross-sectional views of the device, including nanochannels extending beyond the silicon surface to interface with target tissue. **(B)** Experimental design. Sciatic nerve transection was performed to induce muscle denervation, followed by TNT-mediated delivery of *ABM* or sham treatment to the gastrocnemius muscle. Functional assessments were performed weekly for 5 weeks, with endpoint tissue analyses conducted at study completion. Healthy non-denervated mice served as controls. **(C)** Quantitative gene expression analysis of denervated gastrocnemius muscle 24 h after TNT treatment confirming successful delivery and expression of *ABM* transgenes relative to sham-treated controls. **(D)** Kaplan–Meier analysis of needle electromyography (EMG) recordings demonstrating accelerated resolution of denervation-associated fibrillation potentials in *ABM*-treated animals compared with sham controls. **(E–F)** Muscle contractility measurements showing longitudinal assessment of **(E)** twitch torque and **(F)** tetanic torque following denervation and treatment. *ABM*-treated muscles exhibited trends toward improved functional recovery compared with sham-treated controls over the 5-week observation period. (n=5/group) All error bars are shown as SEM. p-value<0.05. One-way ANOVA and two-way ANOVA, as appropriate. Created in BioRender. Salazar, A. (2026).

We hypothesized that activation of neurogenic transcriptional programs in host muscle cells would establish a myoprotective environment during prolonged denervation. To test this hypothesis, a mouse model of muscle denervation was employed, followed by TNT-mediated delivery of the *ABM* factors into the gastrocnemius muscle. Sham-TNT and non-injured mice were included as controls. Functional outcomes were assessed over a 5-week period using electromyography (EMG), electrophysiological analyses, and muscle contractility measurements **(Figure 4B)**.

To verify TNT-mediated gene delivery, denervated gastrocnemius muscles were harvested 24 hours after treatment and analyzed for transgene expression. These analyses confirmed successful expression of the *ABM* factors in TNT-treated muscles relative to sham-treated controls **(Figure 4C)**. Functional measurements were performed at baseline prior to surgery and subsequently weekly intervals for 5 weeks. To further assess neuromuscular integrity during denervation, needle EMG was used to monitor spontaneous fibrillation potentials within the gastrocnemius muscle. Denervated muscles receiving *ABM*-TNT exhibited a more rapid resolution of fibrillation activity compared to sham-TNT controls. Kaplan-Meier analysis demonstrated an earlier disappearance of fibrillation potentials in the *ABM*-treated group, with a marked reduction by week 3 and complete resolution in all treated animals by week 5. In contrast, most sham-treated animals continued to exhibit fibrillation activity at both weeks 3 and 5 **(Figure 4D)**.

Muscle contractile performance was subsequently evaluated using the mPhys system by measuring peak plantar-flexion torque during twitch and tetanic contractions elicited through in vivo sciatic nerve stimulation. Although no statistically significant differences were observed, *ABM*-treated animals exhibited a trend toward improved recovery of both twitch and tetanic torque over the 5-week observation period relative to sham-TNT controls. At week 3, mean twitch torque in the *ABM* group was approximately 2.0-fold higher than sham-TNT and tetanic torque was approximately 2.4-fold higher, before converging toward more modest differences at later time points **(Figure 4E-F)**. Compound muscle action potential (CMAP) amplitudes, a measure of neuromuscular transmission, revealed a pronounced reduction in in both denervated groups at early time points, followed by partial recovery over the 5-week observation period. Consistent with previous findings, CMAP amplitudes also showed a tendency toward improvement in the *ABM*-treated group, at week 3 mean CMAP amplitudes were 2.5-fold higher in the *ABM* group compared to sham-TNT, despite the absence of statistically significant differences **(Figure S2A)**. At study endpoint, triceps surae muscles from denervated and contralateral limbs were harvested and weighed to assess muscle preservation. As expected, denervated muscles from both treatment groups displayed significant reductions in muscle mass relative to their contralateral counterparts. While no significant differences were observed between *ABM*-TNT and sham-TNT groups, *ABM*-treated muscles retained approximately 75% of contralateral muscle mass compared to 63% in sham-treated animals, consistent with a trend toward greater muscle preservation following ABM treatment **(Figure S2B)**. Collectively, these findings demonstrate the feasibility of TNT-mediated delivery of the *ABM* neurogenic cocktail to denervated skeletal muscle. Notably, *ABM* treatment was associated with accelerated resolution of denervation-induced fibrillation activity, a potential indicator of improved neuromuscular stability. Although improvements in contractile performance, CMAP amplitudes, and muscle mass did not reach statistical significance, the observed trends support further investigation of *ABM*-mediated neurogenic reprogramming as a strategy to preserve muscle function during denervation.

### RNA-seq reveals transcriptional programs associated with myoprotection in denervated muscle following *ABM*-TNT treatment

To determine the transcriptomic effects of TNT-mediated delivery of the neurogenic reprogramming cocktail *ABM in vivo*, RNA-seq was performed on denervated gastrocnemius muscle collected 5 weeks after treatment. Global transcriptomic analyses demonstrated substantial differences between denervated and healthy muscle, with *ABM*-TNT-treated muscles exhibiting a distinct transcriptional profile relative to both sham-TNT and healthy control groups **(Figure S3A)**. Differential expression analysis identified 271 upregulated and 74 downregulated genes in *ABM*-treated muscles compared with sham-TNT controls **(Figure 5A)**, and 653 upregulated and 36 downregulated genes relative to healthy muscle **(Figure 5D)**. Consistent with these findings, unsupervised hierarchical clustering of DEGs revealed clear segregation of *ABM*-treated samples from both sham-treated **(Figure S3B)** and healthy control muscles **(Figure S3C)**. Heatmap analyses further highlighted distinct expression patterns associated with muscle maintenance, regeneration, and tissue remodeling in *ABM*-treated muscles compared with both comparison groups **(Figure 5B,E)**. Among the genes enriched in *ABM*-treated muscles, several were associated with muscle regeneration, neuromuscular organization, and trophic support, including Igf1[57], Myh8[58], Ache[50], Mymk, Mymx[59], and Dll1[60]. GO enrichment analysis revealed significant overrepresentation of biological processes related to extracellular matrix organization, skeletal system development, axonogenesis, and angiogenesis in *ABM*-treated muscles relative to both sham-treated and healthy controls **(Figure 5C,F)**. Similarly, GSEA demonstrated enrichment of pathways involved in developmental signaling, myogenesis, extracellular remodeling, and metabolic adaptation, including Notch signaling, myogenesis, angiogenesis, epithelial-to-mesenchymal transition (EMT), and Hedgehog signaling **(Figure 5G)**.

**Figure 5.**
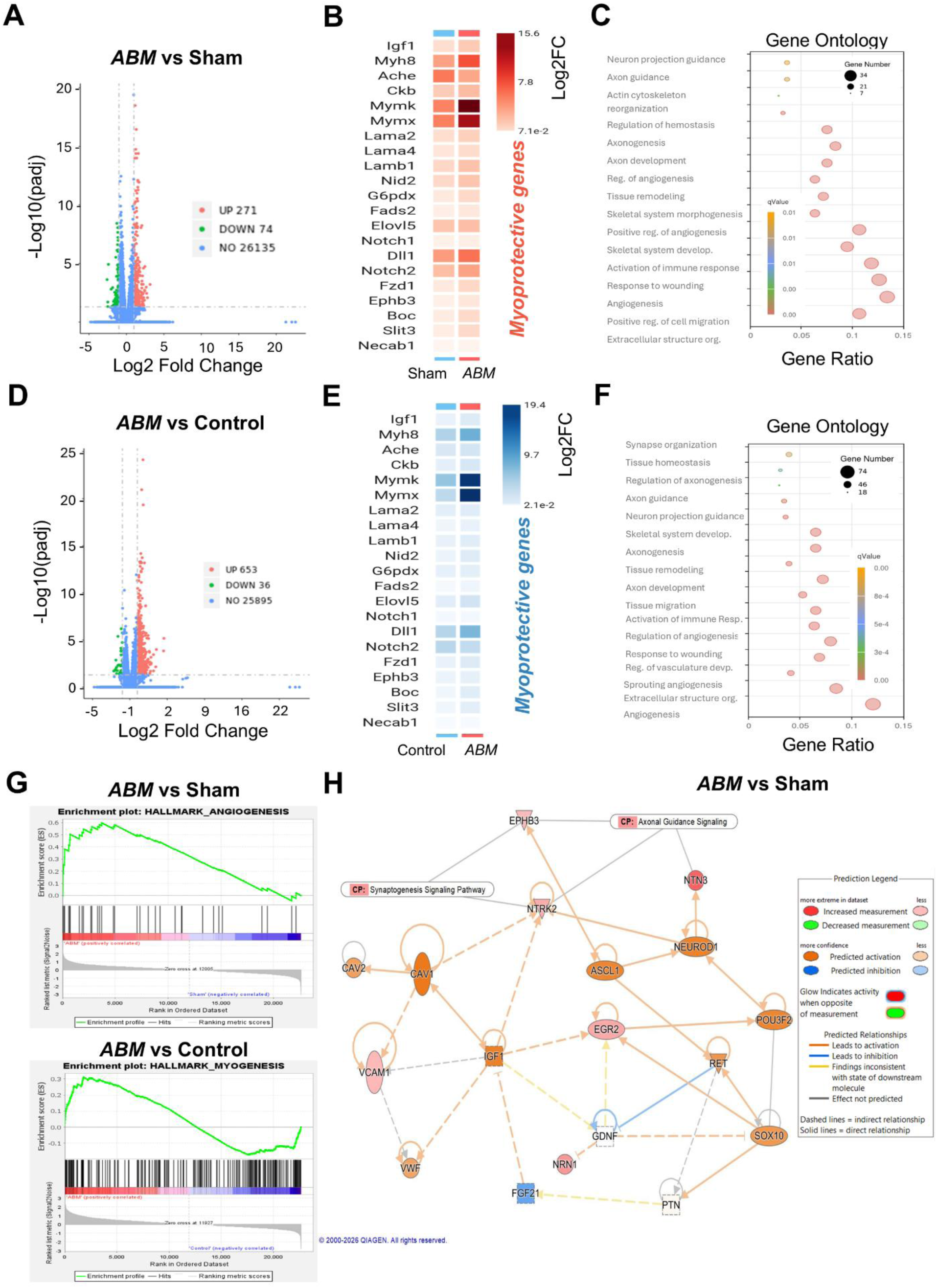
Transcriptomic analyses reveal activation of neurogenic, regenerative, angiogenic, and myoprotective programs in denervated muscle following *ABM* TNT. RNA-seq was performed on denervated gastrocnemius muscles collected 5 weeks after TNT treatment. **(A)** Volcano plot showing DEGs in *ABM*-TNT-treated muscles relative to sham-TNT controls. **(B)** Hierarchical clustering heatmap of selected DEGs associated with muscle regeneration, neuromuscular organization, trophic support, and tissue remodeling in *ABM*-treated muscles compared with sham-TNT controls. Upregulated genes are shown in red, downregulated genes in green, and non-differentially expressed genes in blue. **(C)** GO analysis demonstrating overrepresentation of pathways related to extracellular matrix organization, skeletal system development, axonogenesis, and angiogenesis in *ABM*-treated muscles compared with sham-TNT controls. **(D)** Volcano plot showing DEGs in *ABM*-treated muscles relative to healthy control muscle. **(E)** Hierarchical clustering heatmap of selected DEGs associated with regenerative and myoprotective processes in *ABM*-treated muscles compared with healthy controls. **(F)** GO analysis of *ABM*-treated muscles relative to healthy controls. **(G)** GSEA demonstrating enrichment of pathways associated with myogenesis, angiogenesis, developmental signaling, extracellular remodeling, and metabolic adaptation, including Notch signaling, EMT, and Hedgehog signaling, in *ABM*-treated muscles compared with sham-TNT and healthy controls. **(H)** IPA comparing *ABM*-treated and sham-TNT-treated muscles, highlighting activation of neurogenic transcriptional regulators (ASCL1, POU3F2, NEUROD1, and SOX10), neurotrophic signaling molecules, and vascular- and trophic-support pathways associated with regenerative remodeling. Differentially expressed genes were defined as log2FC ≥ |1| with adjusted p-value ≤ 0.05. IPA network legend is shown on the right.

To further investigate biological networks associated with *ABM* treatment, IPA was performed. Comparison of *ABM*-treated and sham-treated muscles identified a network enriched for neurogenic regulators centered on the transcription factors ASCL1, POU3F2, and NEUROD1, together with the lineage-associated factor SOX10, suggesting activation of early neurogenic programs within the denervated muscle environment[61, 62]. Importantly, this persistent *ABM* transgene expression was not anticipated at 5 weeks, as TNT-mediated delivery relies on episomal plasmids that result in transient expression. Therefore, the observed neurogenic signature likely reflects durable downstream transcriptional reprogramming initiated by *ABM* exposure rather than continued expression of the delivered factors.

In parallel, genes involved in axonal guidance and neurite support, including NTN3, NRN1, and NTRK2, were upregulated, consistent with enhanced neurotrophic signaling[63, 64]. The network also incorporated vascular- and trophic-associated factors, including VCAM1, VWF, and IGF1, suggesting coordinated activation of regenerative, angiogenic, and metabolic support pathways[57] **(Figure 5H)**. Comparison of *ABM*-treated muscles with healthy controls similarly identified activation of neurogenic transcriptional networks. In this analysis, growth factors such as PTN and FGF21 were predicted to be activated, indicating enhanced neurotrophic and metabolic support, while vascular-associated signaling pathways remained prominent within the network **(Figure S3D)**. Collectively, these findings demonstrate that TNT-mediated delivery of *ABM* induces a transcriptional program in denervated skeletal muscle characterized by activation of neurogenic, regenerative, angiogenic, and metabolic pathways that may contribute to enhanced muscle stability and resilience during prolonged denervation.

## CONCLUSION

In summary, this study demonstrates that TNT-mediated non-viral delivery of the neurogenic transcription factor cocktail *ABM* induces sustained neurogenic programs that promote myoprotective responses in denervated skeletal muscle. *ABM* delivery promoted neuronal-like transcriptional and morphological changes within myoblasts in vitro, upregulated neuronal markers as early as 7 days post-transfection with sustained expression through 21 days, and exhibited membrane excitability in a subset of *ABM*-treated myoblasts, while activating broad neurogenic regulatory networks associated with neuronal fate commitment, neuronal differentiation, synaptogenesis, and axon guidance. In a mouse model of skeletal muscle denervation, *ABM*-TNT treatment resulted in accelerated resolution of denervation-associated fibrillation activity and transcriptional activation of regenerative, neuromuscular organizational, trophic support, extracellular matrix remodeling, angiogenic, and metabolic pathways 5 weeks after treatment. Collectively, these findings establish TNT-mediated *ABM* delivery as a minimally invasive, non-viral platform capable of promoting a pro-regenerative and myoprotective tissue environment during prolonged denervation, with potential utility for preserving muscle viability during the time necessary for peripheral nerve regeneration and target muscle reinnervation.

## MATERIALS AND METHODS

### Plasmid preparation

Plasmids were expanded through inoculation in *E.coli* and purified following the manufacturers protocol for plasmid isolation (ZymoPURE II Plasmid Midiprep Kit, cat. no. D4201). Plasmid concentrations were verified using a Nanodrop 2000c Spectrophotometer (ThermoFisher Scientific). DNA plasmids used in this study are listed in Table 1. *ABM* plasmids were a gift from Connie Cepko.

**Table 1.**
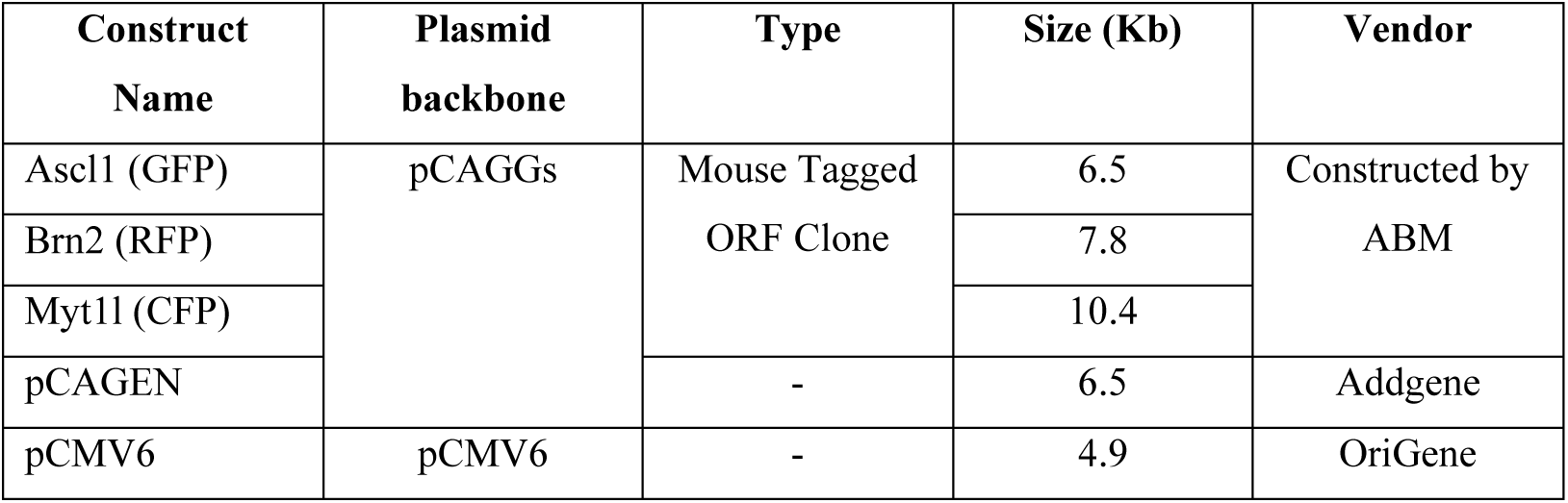
Plasmid information. This table lists all plasmids used in this study, including construct name, Tags, plasmid backbone, type, size (kb), and vendor.

### Cell culture and in vitro non-viral transfection

C2C12s (CRL-1772, ATCC) were seeded and supplemented in Dulbecco’s Modified Eagle Medium (DMEM) (Thermo Fisher Scientific) with 10% fetal bovine serum (FBS, VWR) and 1% penicillin/streptomycin (GIBCO) at 37 °C in a humidified air chamber containing 5% CO_2_. To prevent premature differentiation into myotubes and maintain undifferentiated cell population, confluency was kept below 70%. Once it reached the desired confluency, they were detached, and 1.0 x 10^6^ cells were resuspended in 100 µl of electrolytic buffer for subsequent non-viral cell transfection using a Neon transfection system (Thermo Fisher Scientific). *ABM* plasmids were co-transfected at a 1:1:1 ratio with a concentration of 0.05 μg/μl. Cells were transfected using one 30-ms pulse at 1425 V. After transfection, cells were seeded into 6-well plates and maintained for 24 hours to 21 days, depending on experimental requirements. The media was changed for induction media prepared as follows: DMEM, with 10% FBS, 1% penicillin/streptomycin, 1% N2 supplement (17502048, Thermo Fisher scientific) and 10ng/ml of fibroblast growth factor basic (FGFb, PMG0033, Gibco). C2C12 cultures were transfected with pCMV6 or pCAGEN (sham/only vector) plasmids at a concentration of 0.15 μg/μl in an identical manner as described above to serve as control. Plasmid delivery efficiency of each transcription factor was evaluated by qRT-PCR and by immunofluorescence detection of GFP (Ascl1), RFP (Brn2), and CFP (Myt1l).

### Gene expression assays

Total RNA was extracted using TRIzol Reagent (Invitrogen) according to the manufacturer’s protocol. RNA samples were treated with DNase I (Thermo Fisher) to remove residual DNA, quantified by NanoDrop 2000 spectrophotometry (Thermo Fisher Scientific), and reverse-transcribed using the Maxima First Strand cDNA Synthesis Kit (Thermo Fisher) with 500–2500 ng of RNA in a 20 µl reaction, ensuring equal cDNA input across samples. Gene expression was assessed by qRT-PCR using SYBR Green Master Mix (Thermo Fisher Scientific) with pre-designed probes (Table 2) and TaqMan Fast Advanced Master Mix (Thermo Fisher Scientific) and TaqMan assays (Table 3), with GAPDH serving as the endogenous control.

**Table 2.**
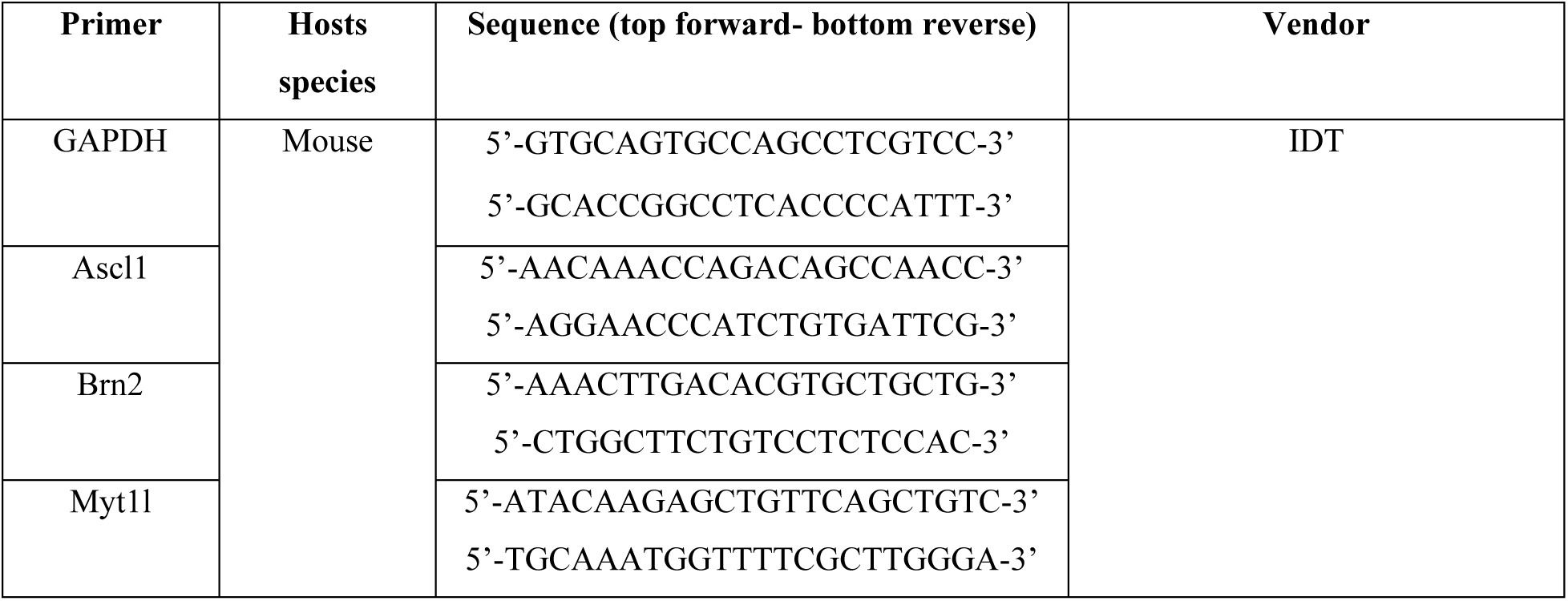
IDT Primer information. This table lists all IDT pre-designed primers used in this study, including the primer name, host species, sequence, and vendor.

**Table 3.** TaqMan Primer information. This table lists all TaqMan pre-design primers used in this study, including the primer name, host species, Assay ID, and vendor.

| Primer | Hosts species | Assay ID | Vendor |
| --- | --- | --- | --- |
| GAPDH | Mouse | Mm99999915_g1 | ThermoFisher Scientific |
| Tuj1 |  | Mm00727586_s1 |  |
| MAP2 |  | Mm00485231_m1 |  |
| Synaptophysin (Syp) |  | Mm00436850_m1 |  |

### Electrophysiology

Whole-cell patch clamp recording was used to measure excitability in bipolar myoblast-derived cells. For recording, cells were continuously superfused with an extracellular bath solution containing 115 mM NaCl, 2 mM KCl, 1.5 mM MgCl2, 3 mM CaCl2, 10 mM HEPES, and 10 mM Glucose (pH 7.4). Glass electrodes (3-4 Mohms) were filled with a pipette solution containing 115 mM K-gluconate, 10mM N-2-Hydroxyethylpiperazine-N’-2-Ethanesulfonic Acid (HEPES), 4 mM NaCl, 0.5 mM ethylene glycol tetraacetic acid (EGTA), 1.5 mM MgCl2, (pH 7.3). Cells had a patch resistance of >100 MOhm after whole-cell access was gained, and series resistance was compensated 40-50%. Data were collected using an Multiclamp 700B amplifier, Digidata 1500B digitizer, and Clampex 11 software (Molecular Devices, San Jose, CA). For analysis of voltage-gated currents, the basal holding potential was -70 mV and cells were stepped for 400 ms in 10 mV increments from -120 mV to 80 mV. Transient inward currents, due to activity of voltage-gated sodium channels, were isolated from measuring the peak amplitude. Sustained plateau currents, reflective of voltage-gated potassium currents, were measured as the average of the last 50ms of the voltage step in the plateau phase of the current. Action potential induction was measured using current clamp. Current was held at 0 pA and then stepped in 3 0 pA intervals for 1 sec.

### TNT Fabrication

TNT devices were fabricated as described by Gallego-Perez et al. [29]. Briefly, projection lithography followed by deep reactive ion etching (DRIE) was used to drill ∼500 nm diameter nanochannels (∼10–15 µm deep) through a 200 µm thick double-side polished silicon (Si) wafer. TNT chips were fabricated using 4”, 200 µm thick, double-side polished silicon wafers in a Class 100 cleanroom. Wafers were first vapor-primed with hexamethyldisilazane (HMDS) to improve photoresist adhesion before spin-coating with AZ1512 positive photoresist (∼1 µm thick). Nano-channels (∼0.8–1 µm dia., 12–15 µm depth) were patterned using direct-write lithography and etched via DRIE. The photoresist was then stripped with acetone. Needles were fabricated on the nano-channel side by patterning and etching microscale pillars. On the wafer’s backside, micro-channels (∼150 µm dia.) were patterned, exposed using lithography, and etched until they connected with the nano-channels. Contact photolithography and DRIE were then used to etch the backside of the wafers to create fluidic access to the nanochannels. Finally, a ∼50 nm-thick insulating layer of silicon nitride was deposited on the surface of the processed silicon using chemical vapor deposition. The devices were also characterized by taking scanning electron microscopy (SEM) images of the cross-section to ensure quality. The wafers were then diced into ∼1 cm² dies, and plasmid reservoirs were created for each die by affixing a plastic ring to the backside of the silicon.

### Animal husbandry

All animal procedures were approved by the Animal Care and Use Committee of The Ohio State University (2016A00000074-R1). C57BL/6J mice were purchased from Jackson Laboratory. Mice were 8-10 weeks at the time of experimentation. Both male and female sexed mice were included in the studies. The animals were housed in groups of no more than five per cage in a sterile, barrier-controlled vivarium under a 12-hour light/dark cycle, with unlimited access to food and water. All mice were acclimated to the housing environment for one week before behavioral assessments and surgical procedures. For postoperative analgesia, mice received Buprenorphine ER (0.1 mg/kg, subcutaneous) before surgery. All animals were anesthetized via isoflurane inhalation before surgery. Anesthesia was induced with 5% isoflurane and maintained with 1.2–2% throughout the procedure. The right hindlimb was shaved and sterilized with three alternating scrubs of 70% isopropyl alcohol and povidone-iodine. A skin incision of ∼2cm was made in the medial aspect of the right leg to expose the sciatic nerve. Sciatic nerves were separated from the surrounding tissue and fascia using Vannas spring scissors. The simple nerve transection injury was performed using the Vannas spring scissors to bisect the nerve. Proximal and distal ends of the nerve were left free-floating. For TNT implementation, blunt dissection was performed to reach the gastrocnemius muscle through the same incision. Then, the TNT chip reservoir containing the plasmid solution (at a 0.05 μg/μl per plasmid concentration) served as the negative electrode. A positive needle electrode was then inserted below the surface of the gastrocnemius, and a pulsed electric field (250V, 10 ms pulses, 10 pulses) was applied across electrodes to drive plasmid DNA across the nanochannels and into the muscle. Once the surgeries were done, the incision was closed with interrupted nylon sutures, and mice were housed individually with mash and hydrogel and placed on heating pad at 37 °C for 2 hours. Upon placement in the vivarium, they were checked for 5 days for postoperative recovery confirmation.

### Functional outcomes

For functional outcomes, baseline measurements were obtained prior to surgeries and over the course of the 5-week period of post-surgery. We employed various electromyography techniques including fibrillations, compound muscle action potential (CMAP), twitch, and tetanic torque. At each session, animals were weighed, and all assessments were performed by the same experimenter to reduce variability, and data collection was blinded to group assignments.

Mice were anesthetized for fibrillations, electrophysiology, and contractility testing by inhaled isoflurane (5% induction, 1.5–2% maintenance) delivered in compressed room air. Artificial tear ointment was applied to the eyes. The body temperature was maintained at 37 °C using a heated table or warming pad, and respiration rate was monitored for the duration of each procedure.

CMAPs were recorded from the triceps surae muscle using a Sierra Summit EMG System (Cadwell, Kennewick, WA), as previously described [36–37]. Briefly, we used sterile 28G insulated monopolar EMG needles placed subcutaneously at the proximal thigh at each side of sciatic nerve. Recording is performed with fine ring electrodes (Catalog # 9013S0312, Natus Neurology, Middleton, Wisconsin, USA) positioned over forelimb or hindlimb with ground electrode placed on tail to minimize artifact. Stimulation was delivered using a portable electrodiagnostic system (Cadwell Sierra Summit, Kennewick, WA) with 0.1-ms pulses at 1–10 mA. Stimulus intensity gradually increased from subthreshold levels until small; all-or-none motor unit responses were detected. The intensity was then raised in quantal steps, repeated for 9 additional increments, followed by acquisition of a supramaximal CMAP.

After electrophysiological recordings, plantarflexion torque was assessed using an *in vivo* contractility apparatus, with a force footplate on rotating axis connected to a force detection motor (Model 1300A, Aurora Scientific Inc., Canada), akin to methods described previously[65, 66]. The left hindpaw was taped to the force plate such that the hindpaw dorsum was angled 90° to the anterior tibia. The leg was extended with the knee joint securely clamped, non-invasively, at the femoral condyles, avoiding compression of the fibular nerve at the fibular head. Two disposable 28-gauge insulated monopolar needle electrodes (Natus Neurology, Inc., Middleton, WI, USA) were inserted subcutaneously over the tibial nerve, just posteromedial to the knee. To identify the maximum stimulus intensity, a series of increasing square stimuli (0.2 ms pulse duration) were delivered at 0.5 Hz frequency until the twitch response no longer increased. Maximal twitch torque was then recorded following a single supramaximal stimulus. Next, a 0.2 ms pulse width of supramaximal square stimuli was delivered at 150 Hz frequency to acquire the maximum tetanic contraction torque.

Single-Fiber EMG (SFEMG) was performed to evaluate neuromuscular junction transmission at the single muscle-fiber level. A sterile concentric needle electrode was inserted into the gastrocnemius to record individual muscle fiber action potentials. The stimulating cathode and anode were placed subcutaneously at the proximal thigh/sciatic notch to activate sciatic motor axons at frequencies ranging from 1–50 Hz to assess both pre and post-synaptic function. A disposable surface disk electrode (CareFusion, Middleton, WI) was positioned on the hind limb, tail, or sacrum to serve as ground and reduce artifacts. Jitter and blocking parameters were quantified. Animal body temperature was maintained at 37°C using a thermoregulated heating pad. Each procedure lasted <30 minutes. After procedures, animals were kept on heating surface pad, and they recovered rapidly from anesthesia. Respiration and peripheral perfusion (pinnae and footpad color) were monitored continuously, and veterinary consultation was sought if complications occurred. For data analysis, the severity of spontaneous activity was graded using a semi-quantitative ordinal scale: 0 (no activity), 1 (persistent activity in ≥2 areas), 2 (moderate activity in ≥3 areas), 3 (widespread activity), and 4 (near-continuous discharges). EMG assessments were performed at baseline and weekly for up to 5 weeks post-surgery. Time to resolution of fibrillation activity was defined as the first time point at which the fibrillation score reached 0 and remained absent in subsequent measurements. Time-to-event analysis was performed using the Kaplan–Meier estimator, with animals not reaching a score of 0 by week 5 treated as censored.

### Muscle Mass Analysis

The triceps of surae muscles were excised from both the experimental and contralateral limbs of the mice. The muscles were weighed individually using an analytical scale.

### RNA sequencing

Total RNA from muscle tissue was isolated using TRIzol Reagent (Invitrogen) and tissue was homogenized using poly-T oligo–attached magnetic beads. RNA was purified with the PureLink RNA Mini Kit (Invitrogen), followed by DNase treatment. Agilent 2100 Bioanalyzer with the RNA 6000 Nano Kit was used to assess RNA quality and samples with RIN ≥ 6 were selected for sequencing by Novogene Co. Libraries were made through end repair, A-tailing, adapter ligation, USER digestion, size selection, PCR amplification, and purification. Library concentration and quality were evaluated by Qubit, qPCR, and Bioanalyzer prior to pooling and sequencing on Illumina platforms to generate FASTQ files. Raw reads were filtered to remove adapters and low-quality sequences, and clean reads were aligned to the mouse reference genome using Hisat2 (v2.0.5). Gene expression was quantified with featureCounts (v1.5.0-p3) and reported as FPKM. Differential gene expression analysis was performed using DESeq2 (v1.20.0) with Benjamini–Hochberg adjusted P-values (FDR ≤ 0.05) and a fold-change threshold of 2 (∣log2FC∣≥1). Gene ontology enrichment analysis was conducted using clusterProfiler, and visualizations were generated using Novomagic, Novogene’s analysis platform. Gene set enrichment analysis (GSEA) was performed using the GSEA software (Broad Institute, UC San Diego) with MSigDB gene sets, considering gene sets with FDR < 25% as significant. Ingenuity Pathway Analysis (IPA, Qiagen) was used to perform pathway and network analyses based on curated molecular interactions in the IPA knowledge base. Canonical pathway activity was inferred using IPA z-score predictions, with positive and negative scores indicating predicted activation or inhibition, respectively. The datasets from this study have been deposited in the Gene Expression Omnibus (GEO) repository with the accession code: GSE334940.

### Statistics

Sigma Plot Version 15.0 and GraphPad Prism were used for all statistical analyses. Data were subjected to normality and outlier tests using Shapiro-Wilk test and ROUT method with Q=1%, respectively. All data is graphed using mean ± standard error of the mean (SEM). Normally distributed data were analyzed using one-way ANOVA with post hoc least significant difference test (Fisher LSD method) or two-tailed t-test as appropriate. A sample size of n=3-5 were used for in vitro experiments and n=5-10 for in vivo experiments. Statistical significance level was defined as P< 0.05.

## Author Contributions

Conceptualization: DGP

Methodology: AISP, SK, JTM, CAVM, CVQ, HH, MF, MEF, JPS, FZ, KD, JA, PB, CDW

Investigation: AISP, SK, JTM, CAVM, CVQ, HH, MF, MEF, JPS, FZ, KD, JA, PB, CDW, JW, ILV, CA, AMM, WDA, DGP

Visualization: DGP, AISP

Funding acquisition: DGP, AMM, WDA

Supervision: DGP, AISP, SK, JTM

Writing – original draft: AISP, SK, JTM, DGP

## Data Availability Statement

All data needed to assess and reproduce the results in this manuscript are exhibited in the paper and/or the Supplemental Materials. GEO with the accession code GSE334940 is available for the bulk RNA-sequencing.

