## Supplemental Material for "Tissue nanotransfection-mediated induction of neurogenic programs promotes myoprotective responses in denervated skeletal muscle"

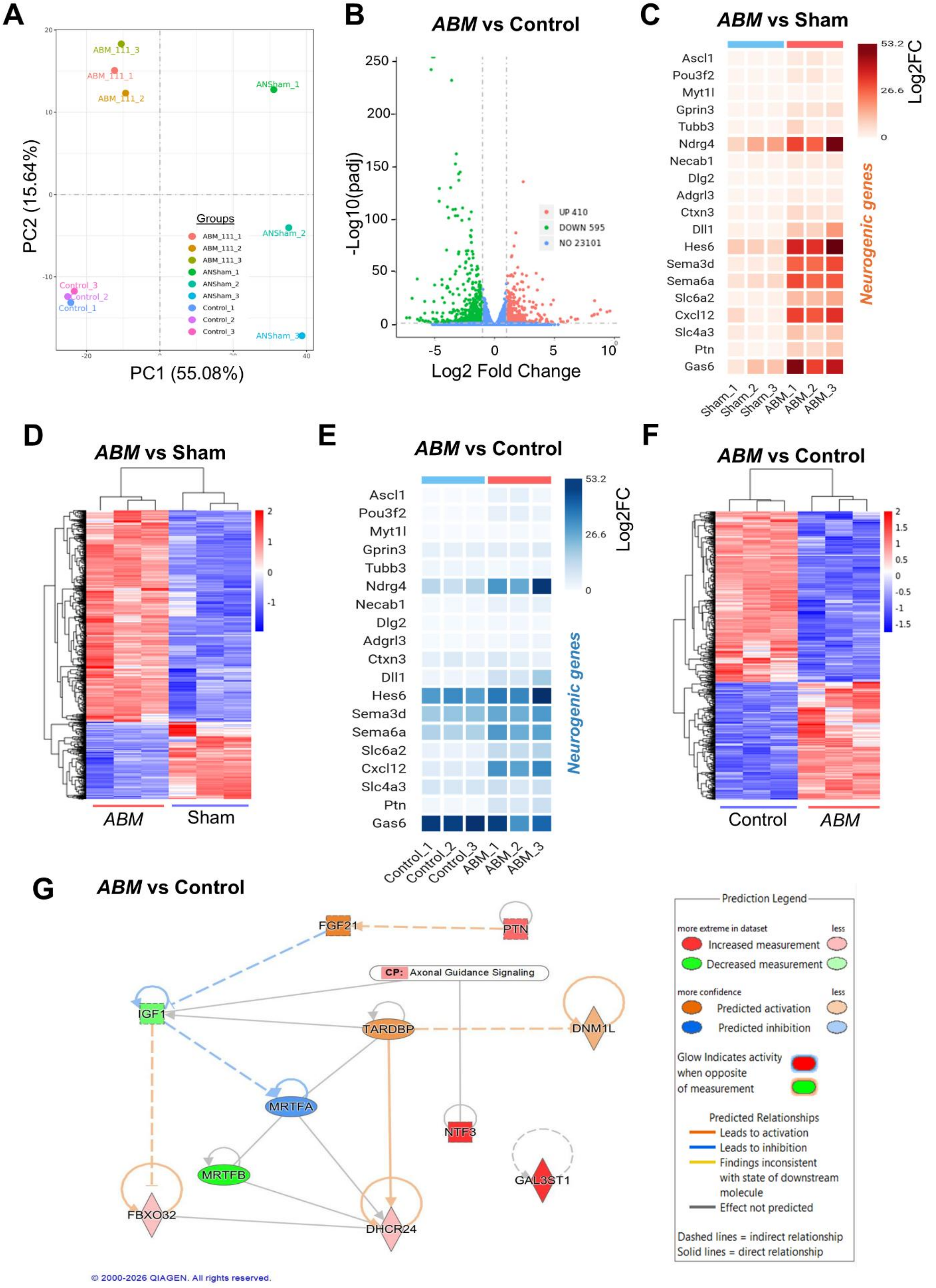
A
PC2 (15.64%)
PC1 (55.08%)
ABM_111_3
ABM_111_1
ABM_111_2
ANSham_1
ANSham_2
ANSham_3
Control_3
Control_2
Control_1
Groups
ABM_111_1
ABM_111_2
ABM_111_3
ANSham_1
ANSham_2
ANSham_3
Control_1
Control_2
Control_3
B
ABM vs Control
-Log10(padj)
Log2 Fold Change
UP 410
DOWN 595
NO 23101
C
ABM vs Sham
Log2FC
53.2
26.6
0
Ascl1
Pou3f2
Myt1l
Gprin3
Tubb3
Ndrg4
Necab1
Dlg2
Adgrl3
Ctxn3
Dll1
Hes6
Sema3d
Sema6a
Slc6a2
Cxcl12
Slc4a3
Ptn
Gas6
Neurogenic genes
Sham_1
Sham_2
Sham_3
ABM_1
ABM_2
ABM_3
D
ABM vs Sham
ABM
Sham
E
ABM vs Control
Log2FC
53.2
26.6
0
Ascl1
Pou3f2
Myt1l
Gprin3
Tubb3
Ndrg4
Necab1
Dlg2
Adgrl3
Ctxn3
Dll1
Hes6
Sema3d
Sema6a
Slc6a2
Cxcl12
Slc4a3
Ptn
Gas6
Neurogenic genes
Control_1
Control_2
Control_3
ABM_1
ABM_2
ABM_3
F
ABM vs Control
Control
ABM
G
ABM vs Control
FGF21
PTN
CP: Axonal Guidance Signaling
IGF1
TARDBP
DNM1L
MRTFA
NTF3
MRTFB
GAL3ST1
FBXO32
DHCR24
© 2000-2026 QIAGEN. All rights reserved.
Prediction Legend
more extreme in dataset
less
Increased measurement
Decreased measurement
more confidence
less
Predicted activation
Predicted inhibition
Glow Indicates activity when opposite of measurement
Predicted Relationships
Leads to activation
Leads to inhibition
Findings inconsistent with state of downstream molecule
Effect not predicted
Dashed lines = indirect relationship
Solid lines = direct relationship

**Figure S1: Transcriptomic changes induced by the *ABM* neurogenic cocktail in myoblasts:** RNA-seq analysis was performed on myoblast cultures 14 days after electrotransfection with *ABM* and compared to sham-transfected cells and non-transfected (control) cells. **(A)** Principal component analysis (PCA) showing clustering of samples and separation of groups. **(B)** Volcano plot depicting DEGs in *ABM*-treated myoblasts compared to non-transfected control cells. Genes in red are significantly upregulated in the *ABM* group relative to the control, genes in green are downregulated and those in blue are not differentially expressed. **(C)** DEG clustering heatmap highlighting neurogenic gene expression pattern in *ABM*-treated cells compared to sham-transfected cells. **(D)** Heatmaps of DEG illustrating distinct transcriptional profiles between *ABM* and sham-transfected cells. Similarly, **(E)** hierarchical clustering heatmap of DEGs showing the neurogenic gene expression and **(F)** heatmaps of DEG demonstrating clear differences between *ABM*-treated cells compared to control myoblasts. **(G)** IPA network highlighting preferential engagement of neurogenic-associated signaling modules in *ABM*-treated myoblasts compared to controls. IPA network legend shown on the right side. Genes with Log2FC $\geq|1|$ and adjusted p-value $\leq 0.05$ were considered significantly differentially expressed.

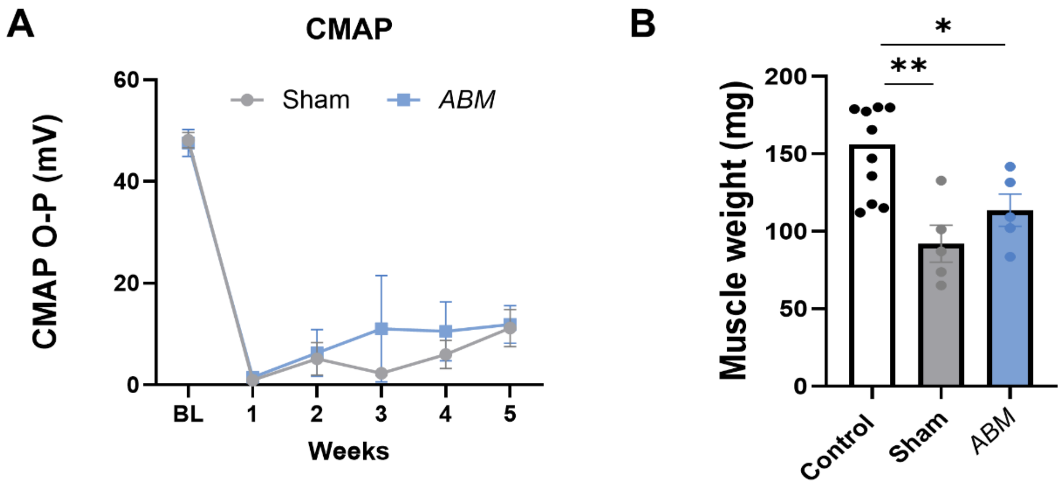


**Figure S2. Functional assessment of denervated muscle following TNT-mediated delivery of *ABM*. (A)** Longitudinal evaluation of CMAP amplitudes following sciatic nerve resection and TNT-mediated delivery of *ABM* or sham treatment to the gastrocnemius muscle. Sciatic nerve stimulation revealed a pronounced reduction in CMAP amplitudes in both denervated groups at early time points, followed by partial recovery over the 5-week observation period. No statistically significant differences were detected between *ABM*-TNT and sham-TNT treated animals. **(B)** Endpoint analysis of gastrocnemius and soleus muscle mass at 5 weeks post-denervation. Both denervated groups exhibited significant muscle atrophy relative to healthy controls. Although differences did not reach statistical significance, muscles receiving *ABM* via TNT demonstrated a trend toward greater preservation of muscle mass compared with sham-TNT controls, with values approaching those of the contralateral non-denervated muscles (n=5/group). All error bars are shown as SEM. p-value<0.05. One-way ANOVA and two-way ANOVA, as appropriate.

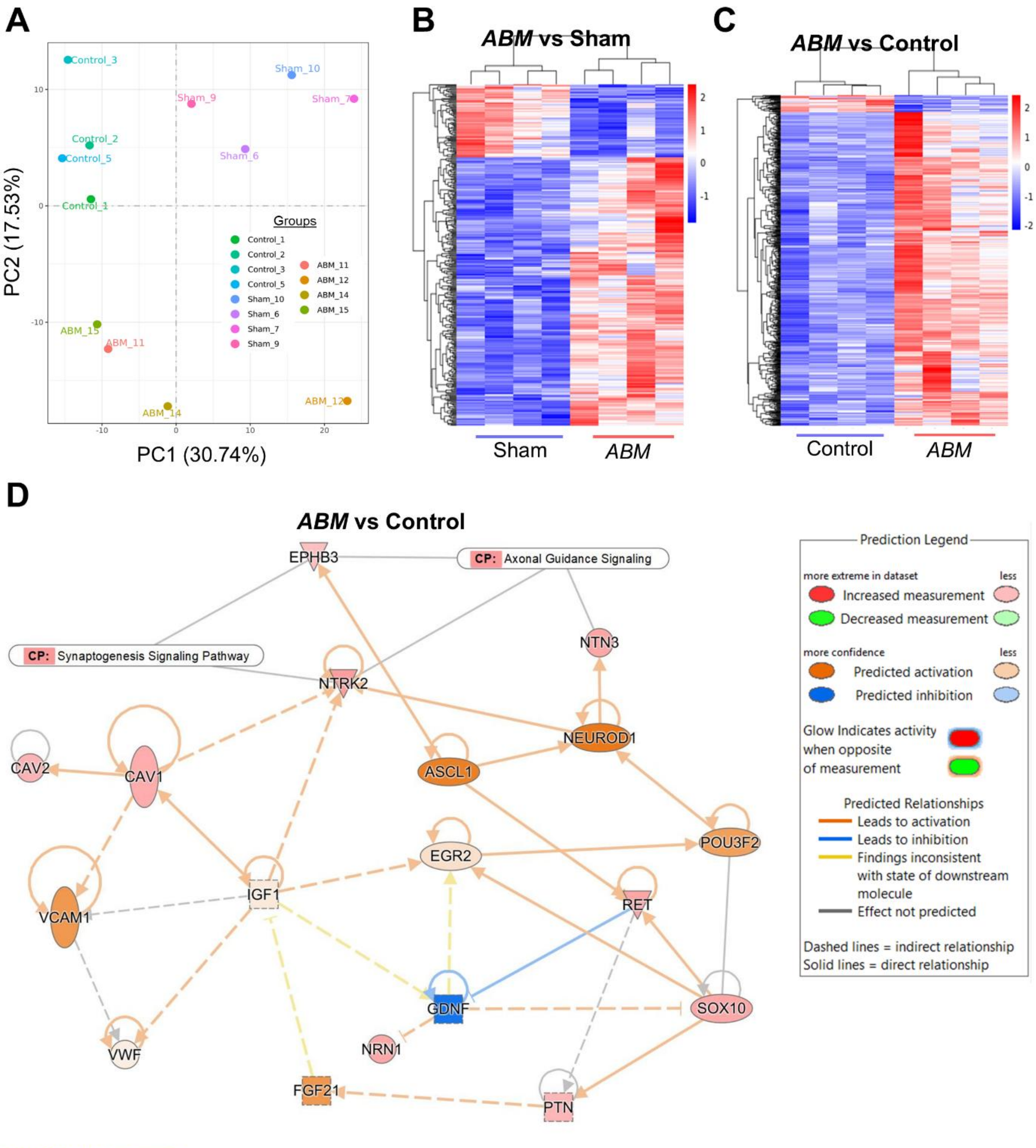


)0-2026 QIAGEN. All rights reserved.

**Figure S3. RNA-seq analysis reveals the impact of neurogenic TNT in denervated muscles.** RNA-seq was performed on denervated gastrocnemius muscles collected 5 weeks after TNT treatment. **(A)** PCA demonstrating distinct clustering of *ABM*-TNT-treated, sham-TNT-treated, and healthy control samples based on global gene expression profiles. **(B,C)** Unsupervised hierarchical clustering of DEGs in *ABM*-TNT-treated muscles compared with **(B)** sham-TNT-treated muscles and **(C)** healthy control muscles, demonstrating distinct transcriptional signatures associated with *ABM*-TNT treatment. DEGs were defined as Log2FC $\geq|2|$ with adjusted p-value $\leq 0.05$. (D) IPA comparing *ABM*-TNT-treated muscles with healthy controls, highlighting activation of neurogenic transcriptional regulators, neurotrophic signaling pathways, and vascular- and metabolic-support networks associated with regenerative remodeling. IPA network legend is shown on the right.
